# mRNA and long non-coding RNA expression profiles in rats reveal inflammatory features in sepsis-associated encephalopathy

**DOI:** 10.1101/117903

**Authors:** WenChong Sun, Ling Pei, Zuodi Liang

## Abstract

**Background:** Sepsis-associated encephalopathy (SAE) is related to cognitive sequelae in patients in the intensive care unit (ICU) and can have serious impacts on quality of life after recovery. Although various pathogenic pathways are involved in SAE development, little is known concerning the global role of long non-coding RNAs (lncRNAs) in SAE.

**Methods:** Herein, we employed transcriptome sequencing approaches to characterize the effects of lipopolysaccharide (LPS) on lncRNA expression patterns in brain tissue isolated from Sprague-Dawley (SD) rats with and without SAE. We performed high-throughput transcriptome sequencing after LPS was intraperitoneally injected and predicted targets and functions using bioinformatics tools. Subsequently, we explored the results in detail according to Gene Ontology (GO) and Kyoto Encyclopedia of Genes and Genomes (KEGG) analyses.

**Results:** LncRNAs were differentially expressed in brain tissue after LPS treatment. After 6 h of LPS exposure, expression of 400 lncRNAs were significantly changed, including an increase in 316 lncRNAs and a decrease in 84 lncRNAs. In addition, 155 mRNAs were differentially expressed, with 84 up-regulated and 71 down-regulated. At 24 h post-treatment, expression of 117 lncRNAs and 57 mRNAs was consistently elevated, while expression of 79 lncRNAs and 21 mRNAs was decreased (change > 1.5-fold; p < 0.05). We demonstrated for the first time that differentially expressed lncRNAs were predicted to be enriched in a post-chaperonin tubulin folding pathway (GO : 007023), which is closely related to the key step in the tubulin folding process.

Interestingly, the predicted pathway (KEGG 04360: axon guidance) was significantly changed under the same conditions. These results reveal that LPS might influence the construction and polarization of microtubules, which exert predominant roles in synaptogenesis and related biofunctions in the rodent central nervous system (CNS).

**Conclusions:** An inventory of LPS-modulated expression profiles from the rodent CNS is an important step toward understanding the function of mRNAs, including lncRNAs, and suggests that microtubule malformation and dysfunction may be involved in SAE pathogenesis.

## Summary

We explored the expression profiles of long non-coding RNAs (lncRNAs) and mRNAs in rodent brain tissues by RNA-sequencing, and identified aberrant expression of 596 lncRNAs and 195 mRNAs ( fold change > 1.5, *P* < 0.05), suggesting lncRNAs were affected by sepsis-associated encephalopathy (SAE). When applying gene ontology and Kyoto Encyclopedia of Genes and Genomes (KEGG) pathway analysis, we discovered that these dysregulated genes were mainly involved in cell shape and synaptic plasticity, such as changes in post-chaperonin tubulin folding. We demonstrated for the first time that NONRATT020317 was differentially expressed in SAE.

## Background

Sepsis is a severe disease with high mortality and morbidity. Although there has been a focus on transcription of the genome, the mechanisms still need to be further addressed, especially when the central nervous system (CNS) is involved. Sepsis-associated encephalopathy (SAE) is the most common complication in septic patients, which manifests mental abnormality and motor dysfunction. This type of diffuse brain dysfunction that occurs secondary to sepsis has limited therapeutic options with poor prognosis. As much as 70% survived septic patients in intensive care unit (ICU) had neurocognitive impairment, such as confusion or coma, even never fully recoverd [1]. To improve the quality of life, alleviate burdens of patients’ families and promote the therapeutic effect on sepsis, we urgently need to clarify the pathogenesis of SAE.

Long non-coding RNAs (lncRNAs) are transcripts >200 nucleotides (nt) that actively participate in genomic regulatory pathways via diverse mechanisms (epigenetic regulation, transcription modulation and post-transcription modulation) [2-4]. LncRNAs were previously viewed as “transcriptional noise” without protein-coding capability [5, 6]. Recently, lncRNAs have been extensively investigated and accumulating evidence indicates that they are emerging as key and essential transcriptional and post-transcriptional mediators in diverse physiological and pathological processes in a tissue-specific manner [7-9]. Nevertheless, lncRNAs are highly expressed in developing and adult mammalian brains [10, 11]. LncRNAs have already been implicated to play key roles in brain development, neural plasticity, cognitive function, and neural organization [10]. In the CNS, their emerging roles in diverse neurodegenerative and neurological disorders have been verified [12-14].

It has been shown recently that aberrant lncRNAs are involved in CNS disturbance in rodents [15], and establishing their functions might help to eliminate the neurological consequences. Consequently, it is reasonable to speculate that lncRNA-directed epigenetic regulation in neural communication is linked to the pathophysiology of cognitive dysfunction. However, despite the abundance and pivotal roles of lncRNAs in brain function, their specific expression pattern and underlying functions in SAE are still poorly understood. To explore the lncRNA landscape in rodent models of CNS endotoxemia, we used RNA-seq to provide insights into the genomic roles of lncRNAs under acute and chronic inflammatory conditions in LPS-induced SAE.

## Methods

### Animals

Eight-week-old Sprague–Dawley rats weighing 150–200 g were approved for use by the Ethics Review Committee for Animal Experimentation of the China Medical University, in compliance with the *Guide for the Care and Use of Laboratory Animals.* The rats were housed under a constant temperature (~23±1°C) with adequate food and water. Rats were given lipopolysaccharide (LPS) to create septic model. Animals were divided into three groups (n=20): normal saline (control); L6, rats were killed 6 h after a single intraperitoneal injection of 5 mg/kg LPS; and L24, rats were euthanized 24 h after a single intraperitoneal injection of 5 mg/kg LPS. Brains were removed and dissected on ice.

### RNA extraction

Total RNA was extracted using TRIzol reagent (Invitrogen, Carlsbad, CA, USA). Total RNA quantity and purity were measured using the Agilent 2100 Bioanalyzer (Agilent Technologies, Santa Clara, CA, USA) and RNA 6000 Nano LabChip Kit (Agilent Technologies). The integrity of the RNA was assessed through agarose gel electrophoresis. RNA samples with RNA integrity number >7.0, OD 260/280 ≥1.8 and OD 260/230 ≥1.5 were viewed as high purity and used in the following experiment.

### Library construction and sequencing

rRNA was removed from 5 μg total RNA (including mRNAs and lncRNAs) using the Ribo-Zero rRNA Removal Kit (Epidemiology version) (Epicenter, Madison, WI, USA). For accuracy and reliability, the raw data were preprocessed by trimming the adaptors and low-quality reads. Following purification, mRNA was fragmented into small pieces using divalent cations under elevated temperature. The cleaved RNA fragments were amplified and reverse-transcribed into the final cDNA library using the mRNA-Seq Sample Preparation Kit (Illumina, San Diego, CA, USA). The average insert size for the paired-end libraries was 300 ± 50 bp. Cluster generation and sequencing were performed on a Hiseq2500 sequencer (Illumina) at LC-BIO (Hangzhou, China) using the paired-end sequencing (100 bp).

### Data analysis

Reads were aligned to the rat genome (from UCSC Genome Browser) using Bowtie 2(2.1.0). The expression level of each transcript in each sample was evaluated by reads per kilobase per million mapped reads (RPKM). RPKM = total exon reads/ [mapped reads (millions) × exon length (kb)]. All the lncRNAs were annotated with the corresponding target genes and potential regulatory mechanisms.

### GO (Gene Ontology) and KEGG (Kyoto Encyclopedia of Genes and Genomes) analyses

RNA-seq provided a new perspective on the coordinated correlation of lncRNA expression in the rodent transcriptome under specific stresses. We investigated the function of all the unknown lncRNAs by data curation and reprocessing. Predicted gene targets were separately analyzed using GO (http://www.geneontology.org/) and KEGG (http://www.kegg.jp/kegg/pathway.html). GO terms were used to depict aberrantly expressed RNAs from three aspects: cellular component, molecular function and biological process. KEGG terms were applied to annotate RNAs in biological pathways. Sequencing analysis was performed by LC-BIO.

### Statistical analysis

Statistical analysis was performed by one-way analysis of variance followed by Student’s *t* test using SPSS version 13.0 (SPSS, Chicago, IL, USA). Quantitative data were expressed as mean ± standard deviation. Statistical significance was noted with a threshold of fold change >1.5 and *p* < 0.05 for analysis of sequencing data, and the false discovery rate was calculated to correct the *p* value. Two-sided Fisher’s exact test was used to designate GO category and KEGG results.

## Results

### Alignment of RNA-seq reads

There were 24,954,481 clean reads and an average of 8,318,160 per sample, with a valid ratio of 99.82%. The unique clean reads were 14,123,311, with a valid ratio of 99.73%. The workflow of sequencing in this study is shown in Fig. 1. The valid clean data were analyzed with Bowtie software, which mapped these data to reference values in the mRNA/lncRNA database for subsequent analysis of differential expression and function of the obtained transcripts (Table 1). The characteristic expression tendency of mRNAs and lncRNAs is shown in Fig. 2.

**Fig 1:**
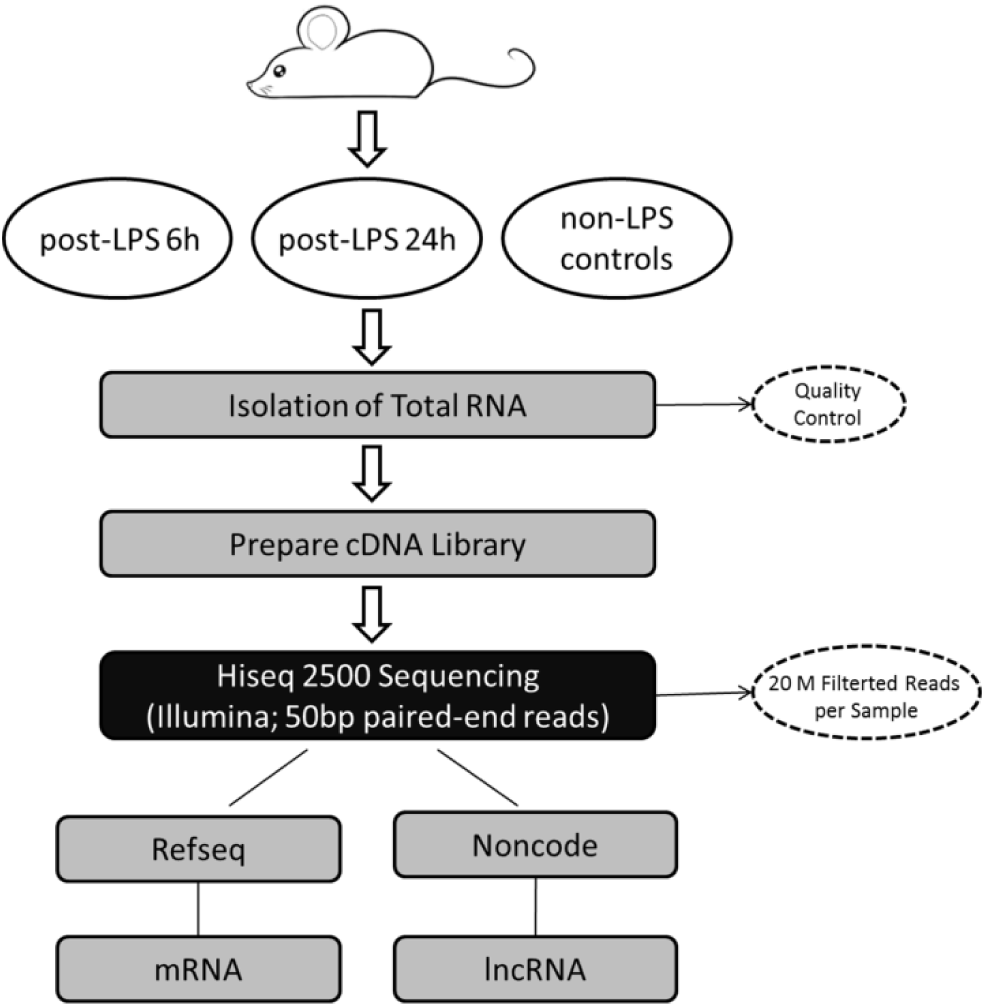
Flowchart of RNA-seq workflow. Schematic representation of analytical procedures for biomarkers of SAE and healthy controls using Hiseq2500 sequencing technology.

**Fig 2:**
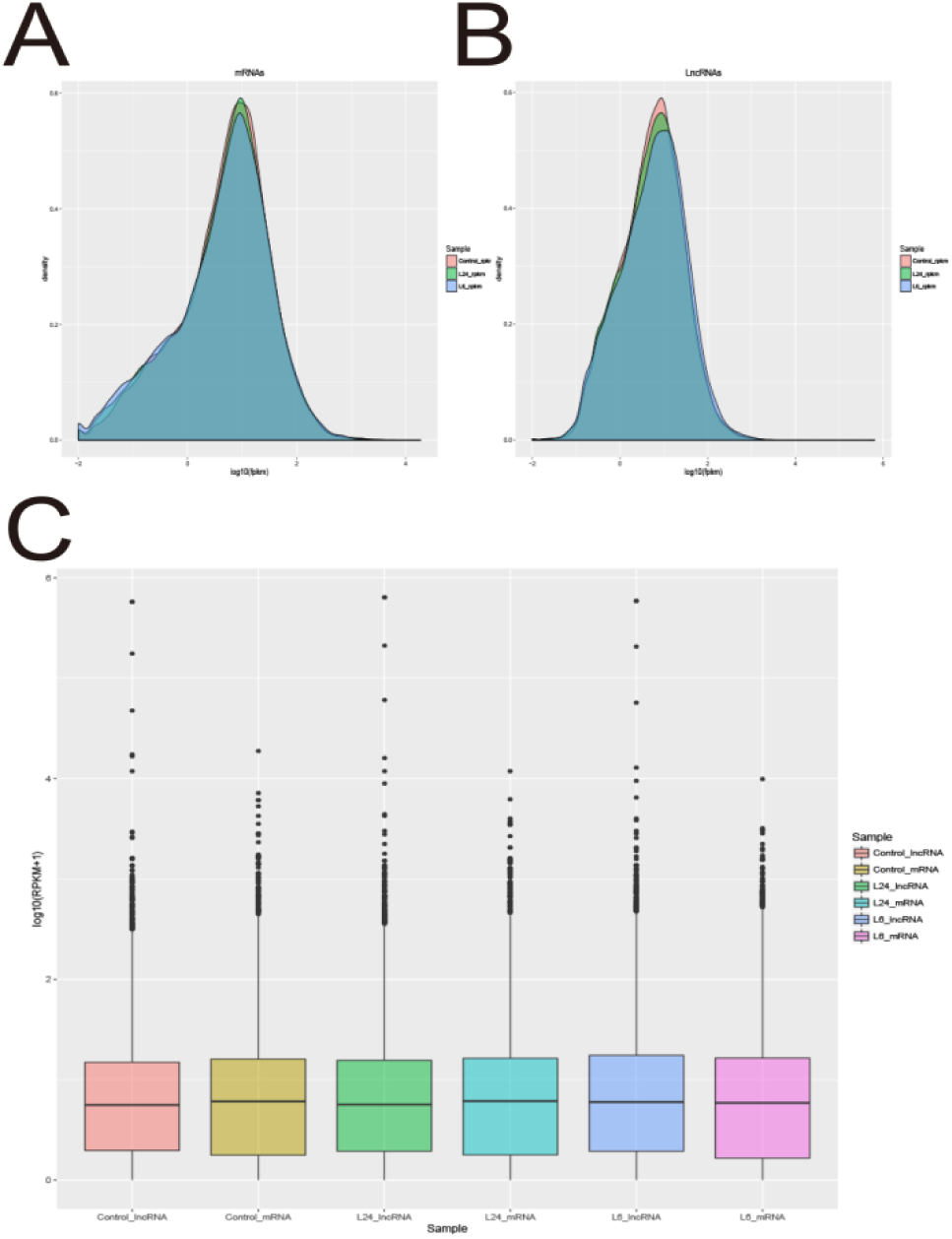
Expression of mRNA and lncRNA in each sample. (A) Distribution of mRNA expression calculated in log_10_ RPKM among three different groups. (B) Distribution of lncRNA expression calculated in log_10_ RPKM among three different groups. (C) Boxplots (showing the 15^th^, 25^th^, 50^th^, 75^th^, and 95^th^ percentiles) show the characteristics of mRNA and lncRNA expression.

**Table 1.** Statistics of manned references of mRNA and IncRNA on valid data

|  | control |  | L6 |  | L24 |  |
| --- | --- | --- | --- | --- | --- | --- |
| Total unique gene |  |  |  |  |  |  |
| mRNA | 30493 | 100% | 30493 | 100% | 30493 | 100% |
| lncRNA | 30832 | 100% | 30832 | 100% | 30832 | 100% |
| Mapped gene |  |  |  |  |  |  |
| mRNA | 24743 | 81.14% | 25037 | 82.11% | 25046 | 82.14% |
| lncRNA | 25730 | 83.45% | 25758 | 83.54% | 25845 | 83.83% |

### Differential expression profile of mRNAs

To reveal the differential expression profile of the RNA transcriptome, the 8-week-old rodent brain mRNAs exposed to 5 mg/kg LPS were identified by sequencing. In addition to the significantly altered genes, Fig. 3A and B shows the changes in mRNA, including the non-significantly expressed genes in the adult rodent brain transcriptome. After short-term exposure to LPS (L6 group), 155 mRNAs were significantly changed, whereas the number decreased to 78 after long duration of LPS exposure (L24 group). Thirty-eight genes were changed in common between the two groups (Fig. 3C). The genes that were significantly up- and down-regulated after LPS treatment are shown in Fig. 3D, as compared with the genes in the control groups. The Hierarchical cluster analysis of the differentially expressed mRNAs in response to the two durations of LPS exposure is shown by the heatmaps in Fig. 3E and F. Integrative analysis of mRNAs, including the fold changes (experimental *vs*. control groups) and *p* values calculated from the normalized expression data, was performed as outlined in Tables 2 and 3. Table 4 gives details of differential expression of mRNAs commonly altered in response to LPS exposure (*p* < 0.05).

**Fig 3:**
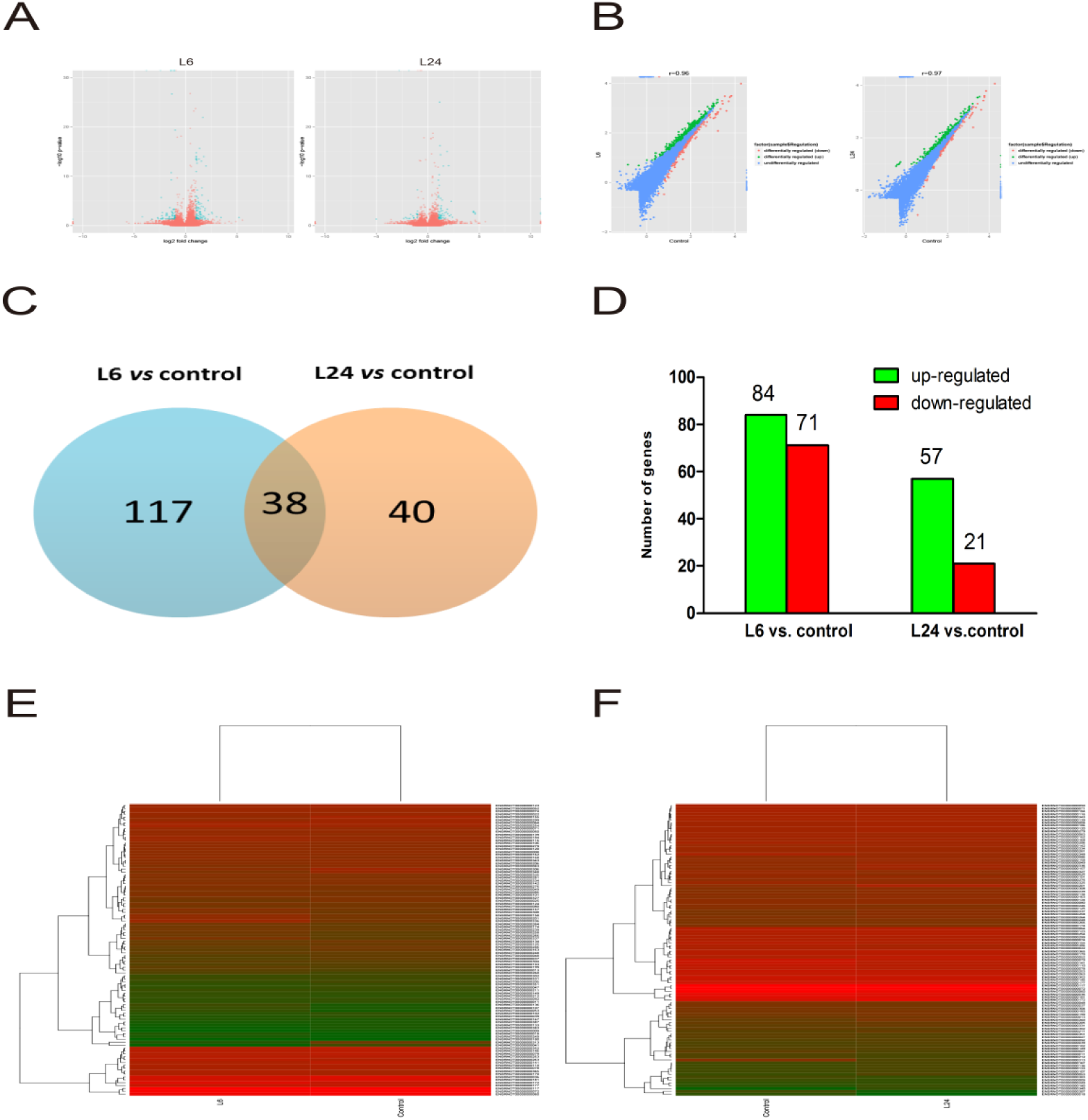
Analysis of differentially expressed mRNAs. (A) Volcano figure showing overall distribution of differentially expressed mRNA. Red indicates the significantly differentially expressed genes (change >1.5-fold and *p* < 0.05) and blue the non-significant ones. (B) Scatter plots give a direct picture of the tendency of data flow, where red indicates the significantly down-regulated genes green the significantly up-regulated genes, and blue plots the non-significant genes. (C) Pie charts illustrate the significantly differentially expressed genes when compared between the L6/L24 and control groups. (D) 141 up-regulated genes (green) and 92 down-regulated genes (red) after different duration of exposure to 5 mg/kg LPS (*p* < 0.05). (E) Hierarchical cluster analysis of mRNA expression in response to 6 h exposure to 5 mg/kg LPS compared to control groups. (F) Hierarchical cluster analysis of mRNA expression in response to 24 h exposure to 5 mg/kg LPS compared to control groups (*p* < 0.05). The clustering tree is listed on the left. Each row represents an individual gene, while colors correspond to different expression variance: green represents lower level, red means higher level, and black means median value (log scale 2, from –2.0 to +2.5).

**Table 2.**
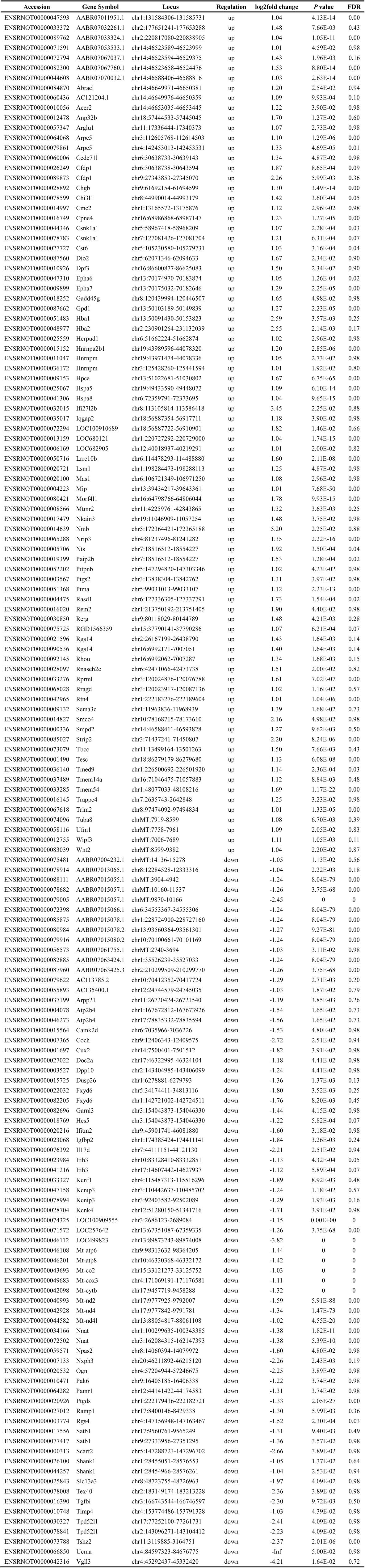
Differentially expressed mRNAs after 6 h exposure

**Table 3.** Differentially expressed mRNAs after 24 h exposure

| Table 3 Differentially expressed mRNAs after 24 h exposure |  |  |  |  |  |  |
| --- | --- | --- | --- | --- | --- | --- |
| Accession | Gene Symbol | Locus | Regulation | log2fold change | P value | FDR |
| ENSRNOT00000085321 | AABR07046866.1 | chr2:41467064-41467252 | up | 1.71 | 3.06E-02 | 0.99 |
| ENSRNOT00000072794 | AABR07067037.1 | chr14:46523658-46524476 | up | 1.22 | 1.22E-02 | 0.99 |
| ENSRNOT00000082300 | AABR07067760.1 | chr5:10125070-10125252 | up | 1.51 | 8.38E-13 | 0.00 |
| ENSRNOT00000060436 | AC121204.1 | chr4:142453013-142453531 | up | 1.30 | 9.53E-05 | 0.03 |
| ENSRNOT00000012478 | Anp32b | chr9:27343853-27345070 | up | 1.94 | 3.85E-03 | 0.56 |
| ENSRNOT00000022943 | Calb2 | chr9:61692154-61694599 | up | 1.45 | 2.55E-02 | 0.99 |
| ENSRNOT00000023869 | Cartpt | chr5:58967418-58968209 | up | 2.38 | 1.97E-02 | 0.99 |
| ENSRNOT00000026249 | Cfdp1 | chr5:62071346-62094633 | up | 1.81 | 1.77E-03 | 0.33 |
| ENSRNOT00000089873 | Cfdp1 | chr4:168752133-168755023 | up | 2.23 | 6.99E-03 | 0.76 |
| ENSRNOT00000078783 | Csnk1a1 | chr8:120439994-120446507 | up | 1.14 | 1.88E-03 | 0.35 |
| ENSRNOT00000012455 | Fuca1 | chr19:41482728-41509658 | up | 1.04 | 2.26E-02 | 0.99 |
| ENSRNOT00000058362 | Gjc2 | chr2:30125249-30127269 | up | 1.18 | 2.14E-02 | 0.99 |
| ENSRNOT00000015152 | Hnrnpa2b1 | chr19:43989596-44078320 | up | 1.23 | 2.99E-06 | 0.00 |
| ENSRNOT00000015325 | Hnrnpa2b1 | chr19:43971474-44078336 | up | 1.02 | 8.85E-05 | 0.03 |
| ENSRNOT00000009153 | Hpca | chr6:72359791-72373695 | up | 1.14 | 9.06E-26 | 0.00 |
| ENSRNOT00000032015 | Ifi27l2b | chr19:55929609-55946250 | up | 4.45 | 1.46E-03 | 0.28 |
| ENSRNOT00000045874 | Igh-1a | chr19:55929664-55946250 | up | Inf | 3.97E-06 | 0.00 |
| ENSRNOT00000031991 | Igkc | chr18:56887722-56910901 | up | 4.59 | 4.41E-03 | 0.59 |
| ENSRNOT00000042165 | Igkc | chr10:16970626-16973418 | up | 4.41 | 1.46E-03 | 0.28 |
| ENSRNOT00000074175 | Igkc | chr5:154269118-154286544 | up | 4.43 | 2.11E-03 | 0.38 |
| ENSRNOT00000086911 | Igkc | chr8:49676540-49703419 | up | 4.59 | 2.53E-03 | 0.43 |
| ENSRNOT00000042900 | LOC100361105 | chr8:49676616-49703410 | up | Inf | 2.53E-03 | 0.43 |
| ENSRNOT00000035257 | LOC680217 | chr10:45526745-45534570 | up | 1.09 | 4.61E-02 | 0.99 |
| ENSRNOT00000006169 | LOC682905 | chr4:81237496-81241282 | up | 1.19 | 6.77E-03 | 0.75 |
| ENSRNOT00000059159 | LOC685203 | chr4:81237496-81241152 | up | 1.23 | 1.46E-02 | 0.99 |
| ENSRNOT00000050716 | Lrrc10b | chr5:147294820-147303346 | up | 1.38 | 6.75E-06 | 0.00 |
| ENSRNOT00000025279 | Mobp | chr6:127336305-127337791 | up | 1.18 | 6.02E-17 | 0.00 |
| ENSRNOT00000025343 | Mobp | chr6:138092131-138093643 | up | 1.02 | 6.22E-10 | 0.00 |
| ENSRNOT00000080421 | Morf4l1 | chr4:98370797-98523273 | up | 1.87 | 9.67E-16 | 0.00 |
| ENSRNOT00000000201 | Mpst | chr4:98337367-98523473 | up | 1.28 | 2.59E-02 | 0.99 |
| ENSRNOT00000008566 | Mtmr2 | chr4:98457810-98523473 | up | 1.28 | 6.99E-03 | 0.76 |
| ENSRNOT00000005706 | Nts | chr9:53440272-53440595 | up | 2.27 | 1.26E-05 | 0.00 |
| ENSRNOT00000019399 | Paip2b | chr10:85573369-85574889 | up | 1.40 | 9.17E-04 | 0.19 |
| ENSRNOT00000066892 | Pitpnb | chr1:226500692-226501920 | up | 1.06 | 3.38E-02 | 0.99 |
| ENSRNOT00000025245 | Plekhb1 | chr2:164628566-164634434 | up | 1.04 | 1.82E-06 | 0.00 |
| ENSRNOT00000082697 | Plekhb1 | chr8:128829680-128842739 | up | 1.04 | 8.87E-07 | 0.00 |
| ENSRNOT00000025093 | Ptma | chr8:128824508-128850358 | up | 1.10 | 4.78E-11 | 0.00 |
| ENSRNOT00000051368 | Ptma | chr8:97474092-97494834 | up | 1.23 | 4.36E-15 | 0.00 |
| ENSRNOT00000004475 | Rasd1 | chr7:119626637-119631425 | up | 1.41 | 3.88E-03 | 0.56 |
| ENSRNOT00000003177 | RGD1310587 | chr8:12284528-12333316 | up | 1.59 | 3.63E-02 | 0.99 |
| ENSRNOT00000066715 | RGD1560608 | chr15:4130808-4132467 | up | Inf | 4.41E-03 | 0.59 |
| ENSRNOT00000021596 | Rgs14 | chr13:93560364-93561301 | up | 1.09 | 2.94E-02 | 0.99 |
| ENSRNOT00000090536 | Rgs14 | chr10:70100661-70101169 | up | 1.05 | 2.94E-02 | 0.99 |
| ENSRNOT00000092145 | Rhou | chr2:121129199-121130675 | up | 1.18 | 9.92E-03 | 0.92 |
| ENSRNOT00000033276 | Rprml | chr17:46322995-46324104 | up | 1.26 | 3.93E-04 | 0.10 |
| ENSRNOT00000015958 | S100a4 | chr10:17048251-17075139 | up | 1.27 | 3.81E-02 | 0.99 |
| ENSRNOT00000023350 | Sord | chr3:154043873-154046330 | up | 1.23 | 3.81E-02 | 0.99 |
| ENSRNOT00000085027 | Strip2 | chr7:44111151-44121130 | up | 1.88 | 4.78E-04 | 0.11 |
| ENSRNOT00000073079 | Tbcc | chr4:115487313-115516296 | up | 1.66 | 2.72E-03 | 0.44 |
| ENSRNOT00000036140 | Tmed9 | chr12:51281987-51341663 | up | 1.03 | 1.42E-03 | 0.28 |
| ENSRNOT00000033285 | Tmem54 | chr1:165665911-165680176 | up | 1.10 | 7.32E-09 | 0.00 |
| ENSRNOT00000022863 | Tppp3 | chr1:165665909-165680206 | up | 1.03 | 1.98E-02 | 0.99 |
| ENSRNOT00000022113 | Ttr | chr9:93545396-93549436 | up | 1.57 | 8.46E-03 | 0.85 |
| ENSRNOT00000074081 | U2af1 | chr13:89873243-89874008 | up | 1.10 | 5.36E-03 | 0.67 |
| ENSRNOT00000058116 | Ufm1 | chr10:46330368-46332172 | up | 1.12 | 2.53E-02 | 0.99 |
| ENSRNOT00000012755 | Wipf3 | chr13:102780877-102790639 | up | 1.26 | 2.76E-04 | 0.07 |
| ENSRNOT00000019860 | Zic1 | chr6:135856218-135861059 | up | 2.09 | 1.07E-02 | 0.97 |
| ENSRNOT00000073231 | AABR07008066.2 | chr17:9777925-9792007 | down | -1.84 | 3.67E-02 | 0.99 |
| ENSRNOT00000079005 | AABR07015057.1 | chr17:9777842-9791781 | down | -1.84 | 0 | 0 |
| ENSRNOT00000056573 | AABR07061755.1 | chr13:88054817-88061108 | down | -1.06 | 2.48E-02 | 0.99 |
| ENSRNOT00000010289 | Apold1 | chr19:56435577-56441733 | down | -1.75 | 3.46E-02 | 0.99 |
| ENSRNOT00000037199 | Arpp21 | chr10:91710495-91711499 | down | -1.01 | 6.99E-03 | 0.76 |
| ENSRNOT00000007365 | Coch | chr2:189997129-189999604 | down | -2.17 | 2.22E-02 | 0.99 |
| ENSRNOT00000068231 | Cpne7 | chr11:87722350-87733734 | down | -1.01 | 1.72E-02 | 0.99 |
| ENSRNOT00000077701 | Cpne7 | chr3:114176309-114207366 | down | -1.04 | 6.10E-03 | 0.72 |
| ENSRNOT00000005383 | Dusp1 | chr4:57204944-57246675 | down | -1.66 | 5.96E-03 | 0.71 |
| ENSRNOT00000022032 | Fxyd6 | chr12:24682041-24710019 | down | -1.46 | 1.17E-02 | 0.99 |
| ENSRNOT00000082205 | Fxyd6 | chr9:16405185-16406338 | down | -1.38 | 2.56E-02 | 0.99 |
| ENSRNOT00000046112 | LOC499823 | chr17:9560761-9565249 | down | -2.79 | 0 | 0 |
| ENSRNOT00000072274 | LOC691153 | chr5:147288723-147296702 | down | -2.03 | 3.46E-02 | 0.99 |
| ENSRNOT00000018402 | Lxn | chr19:37424324-37427989 | down | -1.02 | 5.56E-03 | 0.68 |
| ENSRNOT00000039398 | Neurl1b | chr3:166743544-166746597 | down | -1.38 | 3.02E-02 | 0.99 |
| ENSRNOT00000034166 | Nnat | chr18:15532963-15540177 | down | -1.02 | 5.30E-08 | 0.00 |
| ENSRNOT00000003774 | Rgs4 | chr20:10396621-10407554 | down | -1.00 | 5.56E-03 | 0.68 |
| ENSRNOT00000000313 | Scarf2 | chr17:77252100-77261731 | down | -2.37 | 3.55E-02 | 0.99 |
| ENSRNOT00000049601 | Stx1a | chr2:143096271-143104412 | down | -1.01 | 1.21E-03 | 0.25 |
| ENSRNOT00000073788 | Tshz2 | chr4:84597323-84676775 | down | -1.11 | 3.72E-03 | 0.55 |
| ENSRNOT00000066850 | Ucma | chr8:98734295-98738446 | down | -5.02 | 4.69E-02 | 0.99 |

**Table 4.**
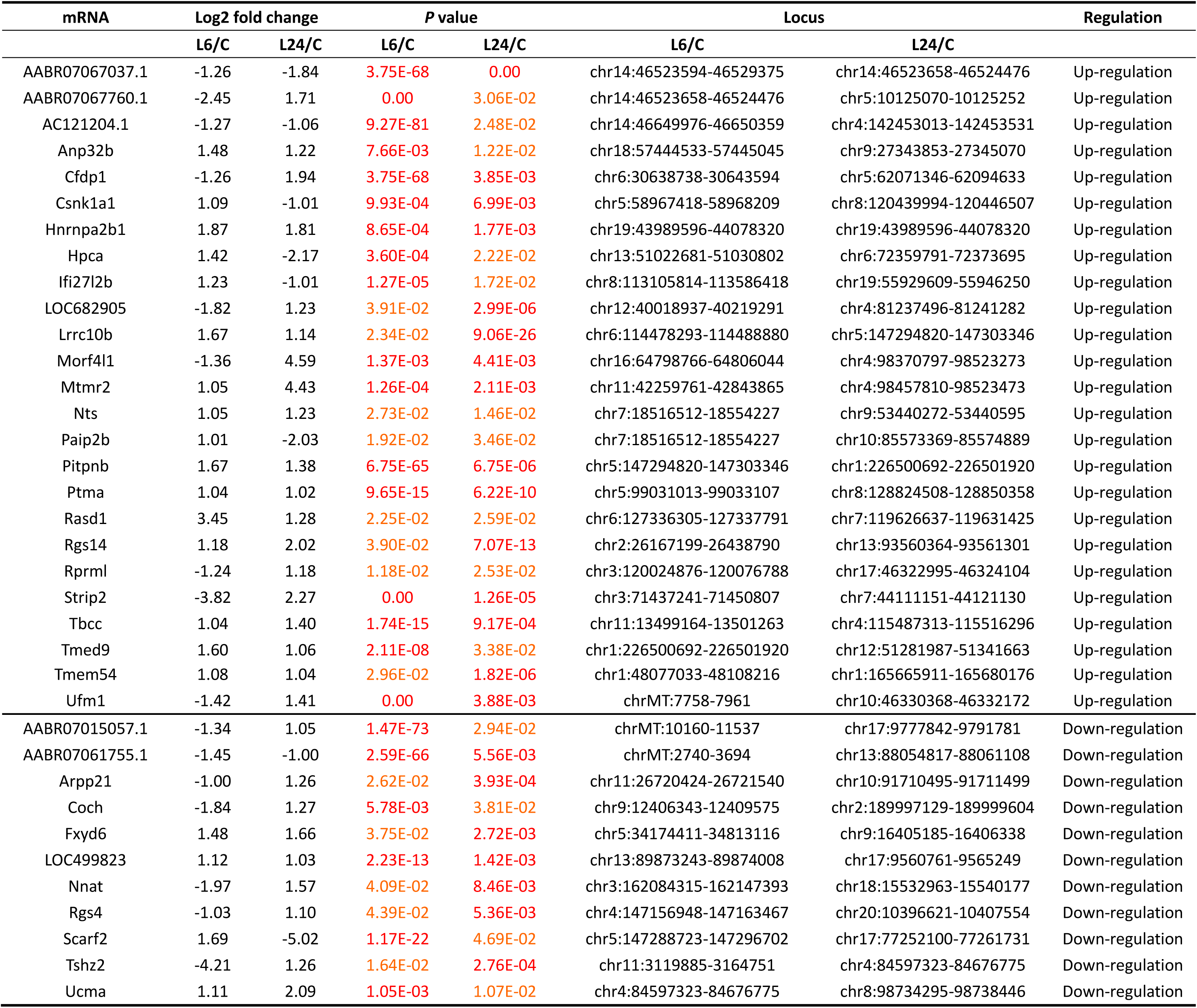
Common differentially expressed mRNAs after 6 h and 24 h exposure

### GO and KEGG analysis

To elucidate further the alterations of cellular component, molecular function, biological process and biological pathways induced by LPS, GO annotations and KEGG analysis were subsequently applied to the differentially expressed genes.

In the L6/C (L6 vs. control) group, the up-regulated mRNAs were involved in 96 biological processes, 16 cellular components, and 29 molecular functions. The down-regulated mRNAs were involved in 85 biological processes, 16 cellular components, and 31 molecular functions. In the L24/C (L24 vs. control) group, the up-regulated mRNAs were involved in 64 biological processes, 8 cellular components, and 35 molecular functions. The down-regulated mRNAs were involved in 22 biological processes, 6 cellular components, and 11 molecular functions. The most frequently represented biological process items were “response to hyperoxia (GO:0055093)” in the L6/C group and “post-chaperonin tubulin folding pathway (GO:007023)” in the L24/C group. The most significant cellular component terms were “respiratory chain (GO:0070469)”, “respiratory chain complex IV (GO:0045277)”, and “mitochondrial respiratory chain complex I (GO:0005747)” in the L6/C group, and “gap junction (GO:0005921)” and “extracellular exosome (GO:0070062)” in the L24/C group. The most frequent molecular function terms were “NADH dehydrogenase (ubiquinone) activity (GO:0008137)”, “GTPase activity (GO:0003924)” and "receptor signaling complex scaffold activity (GO:0030159)” in the L6/C, and “antigen binding (GO:0003823)” in the L24/C groups (Fig. 4A–D).

**Fig 4:**
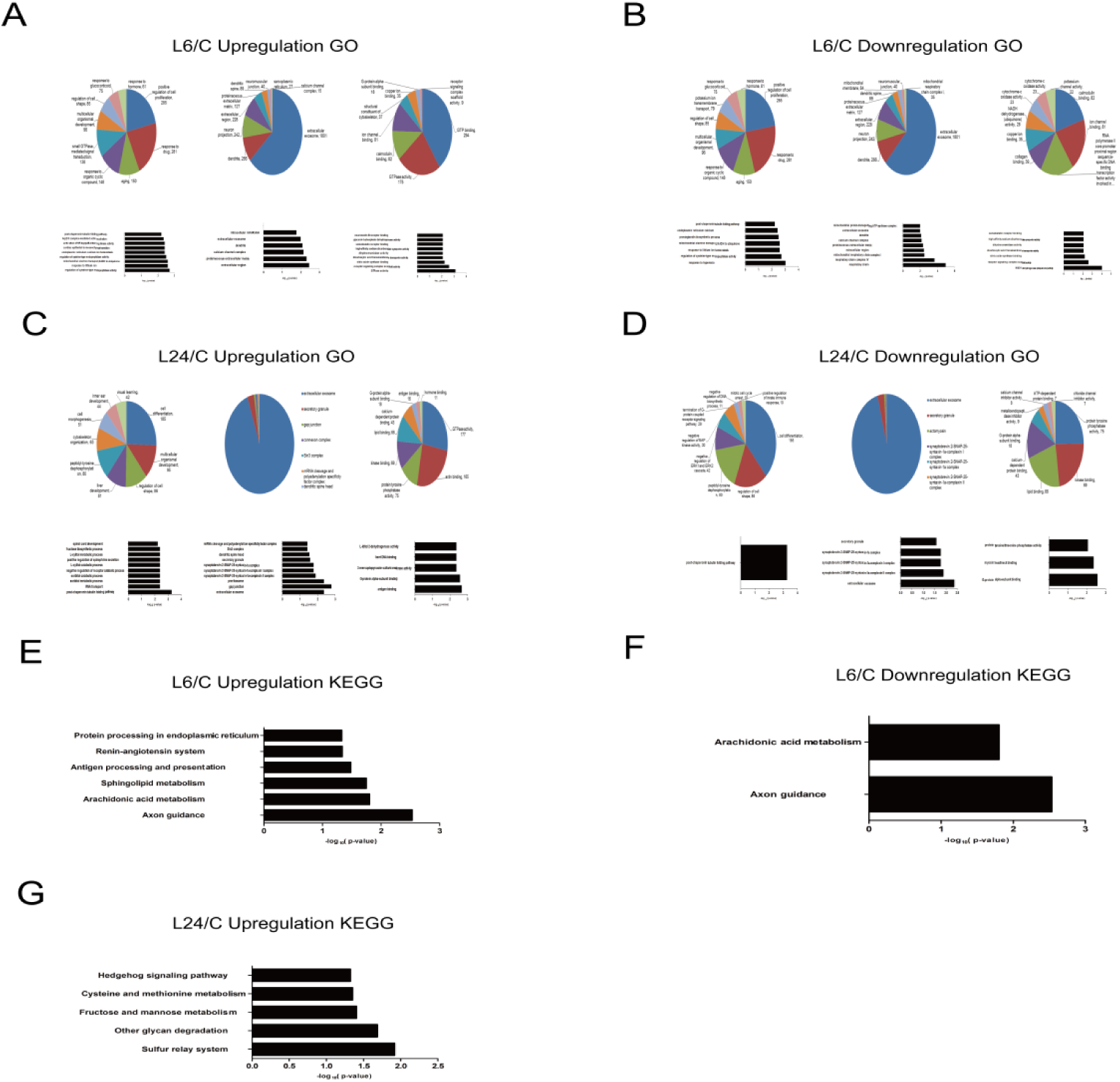
GO and KEGG analysis. (A) Significantly enriched GOs of up-regulated mRNAs in L6/C group. (B) Significantly enriched GOs of down-regulated mRNAs in L6/C group. (C) Significantly up-regulated GOs of up-regulated mRNAs in L24/C group. (D) Significantly down-regulated GOs of down-regulated mRNAs in L24/C group. Results in A–D were all analyzed from the following three aspects of GO analysis: biological process, cellular component, and molecular function. (E) Significantly enriched pathways of up-regulated mRNAs in L6/C group. (F) Significantly enriched pathways of down-regulated mRNAs in L6/C group. (G) Significantly up-regulated pathways of up-regulated mRNAs in L24/C group.

KEGG analysis revealed the substantial enriched pathways of the significantly aberrantly expressed genes (Fig. 4E–G). Compared with the control groups, 6 pathways were significantly up-regulated, and the highest enrichment score in the pathways was “axon guidance” in the L6/C group. The other up-regulated pathways were “arachidonic acid metabolism”, “sphingolipid metabolism”, “antigen processing and presentation”, “renin-angiotensin system” and “protein processing in endoplasmic reticulum” in the L6/C group. Similarly, five pathways were significantly up-regulated in the L24/C group; the top three were “sulfur relay system”, “other glycan degradation” and “fructose and mannose metabolism”.

### Differential expression profile of IncRNAs

Figure 5A and B shows the changes in lncRNAs in the mature rodent brain transcriptome after LPS exposure. Compared with the control groups, we identified 316 significantly differentially expressed up-regulated and 84 significantly differentially expressed down-regulated lncRNAs after short-term exposure to LPS (L6 group) (fold change >1.5, *p* < 0.05). The numbers decreased to 117 and 79 after long-term exposure to LPS (L24 group) (Fig. 5C). Altogether, it revealed that lncRNAs, both up-regulated and down-regulated in L6 group, changed more than with long-term duration. Hierarchical cluster analysis of the differentially expressed lncRNAs is shown by the heatmaps in Fig. 5D and E. The integrative analysis of significantly expressed down-regulated lncRNAs is shown in Table 5.

**Fig 5.**
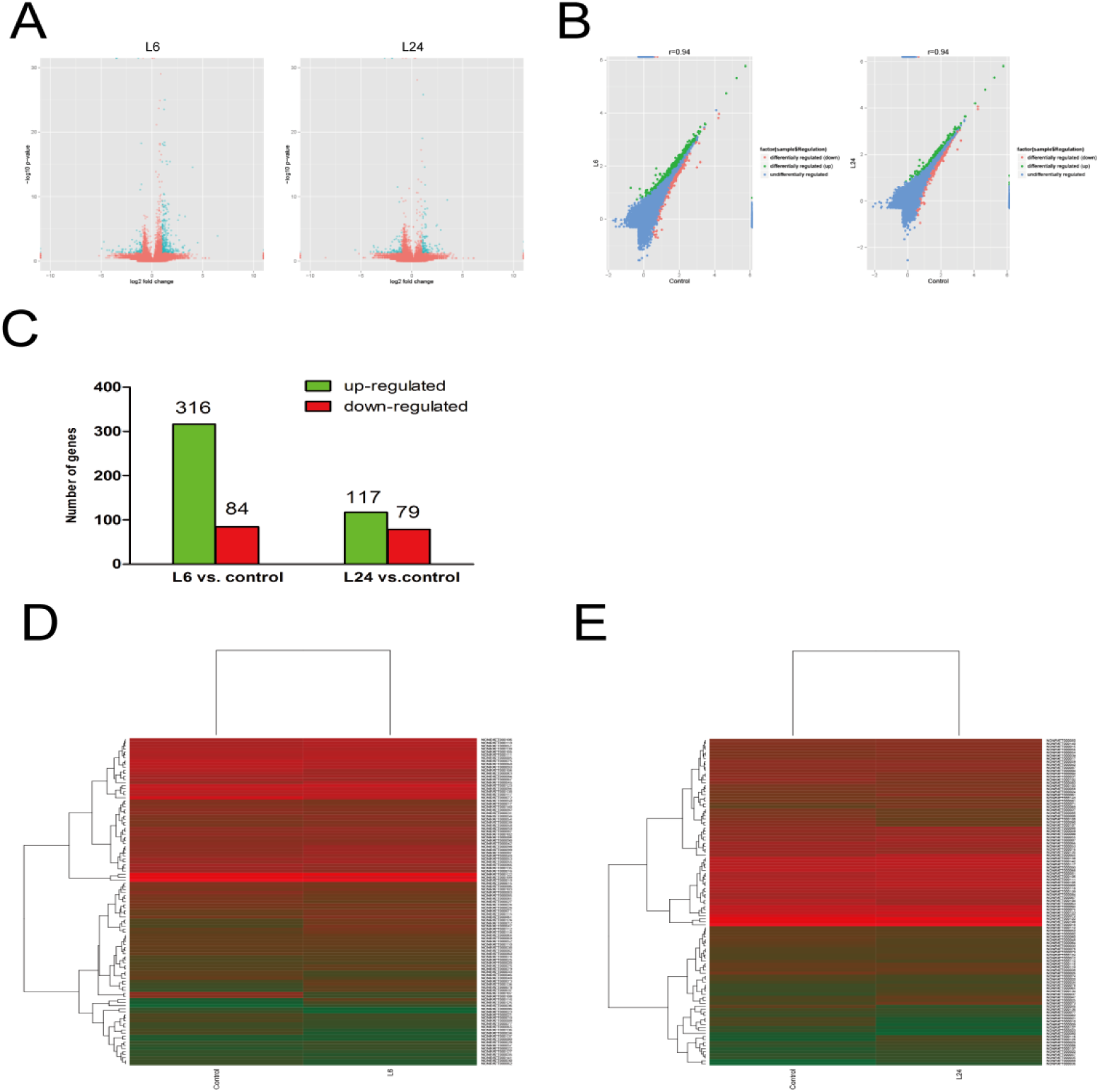
Analysis of differentially expressed IncRNAs. (A), Volcano showing the overall distribution of differentially expressed lncRNAs. Red indicates the significantly differentially expressed lncRNAs (change >1.5-fold and *p* < 0.05) and blue the non-significant ones. (B) Scatter plots give a direct picture of the tendency of data flow, where red indicates the significantly down-regulated lncRNAs, green the significantly up-regulated lncRNAs, and blue means median value. (C) 433 up-regulated lncRNAs (green) and 163 down-regulated lncRNAs (red) after different duration of exposure to 5 mg/kg LPS. (D) Cluster analysis of lncRNA expression in response to 6 h exposure to 5 mg/kg LPS compared to control groups. (E) Cluster analysis of lncRNA expression in response to 24 h exposure to 5 mg/kg LPS compared to control groups (*p* < 0.05). The clustering tree is listed on the left. Each row represents an individual gene, while colors corresponds to different expression variance: green means lower level, red means higher level, and black means the median value (log scale 2, from –2.0 to +2.5).

**Table 5.** Significantly changed down-regulated IncRNAs

| Table 5 Significantly changed down-regulated lncRNAs |  |  |  |  |  |
| --- | --- | --- | --- | --- | --- |
| Transcript id | Log2fold change | P value | FDR | regulation | Transcript type |
| L6/C |  |  |  |  |  |
| NONRATT000107 | -3.26 | 4.46E-03 | 0.20 | down | antisense |
| NONRATT000108 | -2.96 | 1.43E-02 | 0.44 | down | antisense |
| NONRATT000532 | -1.13 | 4.61E-02 | 0.85 | down | sense_overlapping |
| NONRATT001122 | -1.79 | 4.33E-02 | 0.82 | down | sense_overlapping |
| NONRATT001483 | -2.13 | 1.45E-02 | 0.44 | down | antisense |
| NONRATT002002 | -1.24 | 2.56E-02 | 0.63 | down | lincRNA |
| NONRATT002509 | -1.23 | 1.29E-02 | 0.41 | down | sense_overlapping |
| NONRATT002726 | -Inf | 3.77E-02 | 0.77 | down | lincRNA |
| NONRATT003070 | -1.06 | 5.16E-19 | 0.00 | down | sense_overlapping |
| NONRATT003337 | -1.32 | 2.56E-02 | 0.63 | down | lincRNA |
| NONRATT004080 | -1.54 | 3.73E-02 | 0.77 | down | lincRNA |
| NONRATT004414 | -2.29 | 4.62E-02 | 0.85 | down | sense_intronic |
| NONRATT005352 | -4.94 | 1.09E-02 | 0.37 | down | sense_overlapping |
| NONRATT005957 | -1.44 | 4.03E-02 | 0.80 | down | sense_overlapping |
| NONRATT006395 | -4.74 | 1.01E-03 | 0.07 | down | lincRNA |
| NONRATT006508 | -1.59 | 1.91E-02 | 0.52 | down | lincRNA |
| NONRATT006743 | -2.12 | 2.51E-02 | 0.62 | down | lincRNA |
| NONRATT006758 | -2.59 | 1.43E-02 | 0.44 | down | sense_overlapping |
| NONRATT006823 | -1.25 | 3.91E-02 | 0.79 | down | sense_overlapping |
| NONRATT006926 | -1.96 | 4.65E-04 | 0.04 | down | sense_overlapping |
| NONRATT007622 | -3.36 | 3.77E-02 | 0.77 | down | antisense |
| NONRATT008210 | -1.10 | 1.52E-02 | 0.45 | down | lincRNA |
| NONRATT008590 | -3.04 | 3.77E-02 | 0.77 | down | sense_intronic |
| NONRATT009224 | -2.24 | 4.33E-02 | 0.82 | down | lincRNA |
| NONRATT009376 | -1.35 | 0.00E+00 | 0 | down | lincRNA |
| NONRATT009377 | -3.48 | 0.00E+00 | 0 | down | lincRNA |
| NONRATT009710 | -2.94 | 4.62E-02 | 0.85 | down | antisense |
| NONRATT010002 | -1.36 | 3.22E-02 | 0.71 | down | antisense |
| NONRATT010669 | -1.01 | 2.84E-02 | 0.67 | down | antisense |
| NONRATT011133 | -1.19 | 4.97E-02 | 0.88 | down | sense_intronic |
| NONRATT011680 | -2.66 | 4.62E-02 | 0.85 | down | sense_overlapping |
| NONRATT012653 | -1.22 | 2.54E-05 | 0.00 | down | sense_intronic |
| NONRATT012759 | -2.17 | 4.27E-03 | 0.19 | down | sense_overlapping |
| NONRATT013156 | -1.81 | 4.33E-02 | 0.82 | down | antisense |
| NONRATT015238 | -1.01 | 4.37E-03 | 0.19 | down | lincRNA |
| NONRATT015617 | -1.06 | 1.92E-02 | 0.52 | down | lincRNA |
| NONRATT015764 | -1.47 | 1.91E-03 | 0.11 | down | lincRNA |
| NONRATT015765 | -1.25 | 2.56E-02 | 0.63 | down | lincRNA |
| NONRATT015907 | -1.15 | 3.91E-02 | 0.79 | down | sense_overlapping |
| NONRATT017287 | -2.21 | 4.62E-02 | 0.85 | down | sense_overlapping |
| NONRATT017495 | -1.46 | 2.55E-02 | 0.63 | down | sense_overlapping |
| NONRATT018202 | -1.02 | 2.23E-02 | 0.58 | down | lincRNA |
| NONRATT018321 | -3.49 | 0.00E+00 | 0 | down | sense_overlapping |
| NONRATT018471 | -1.02 | 3.61E-02 | 0.76 | down | sense_overlapping |
| NONRATT018581 | -1.09 | 1.28E-04 | 0.01 | down | sense_overlapping |
| NONRATT018620 | -2.91 | 2.56E-02 | 0.63 | down | lincRNA |
| NONRATT018926 | -1.70 | 1.07E-03 | 0.07 | down | sense_overlapping |
| NONRATT019181 | -2.84 | 1.43E-02 | 0.44 | down | sense_overlapping |
| NONRATT019524 | -1.15 | 2.64E-03 | 0.13 | down | lincRNA |
| NONRATT019894 | -1.05 | 1.22E-12 | 0.00 | down | lincRNA |
| NONRATT019895 | -1.15 | 8.08E-12 | 0.00 | down | lincRNA |
| NONRATT019912 | -1.48 | 3.11E-02 | 0.70 | down | lincRNA |
| NONRATT020004 | -1.39 | 3.22E-02 | 0.71 | down | antisense |
| NONRATT020688 | -1.52 | 1.07E-02 | 0.37 | down | sense_overlapping |
| NONRATT020689 | -1.53 | 6.53E-04 | 0.05 | down | lincRNA |
| NONRATT021678 | -2.94 | 3.77E-02 | 0.77 | down | sense_overlapping |
| NONRATT022314 | -3.84 | 2.01E-02 | 0.54 | down | sense_overlapping |
| NONRATT022337 | -3.48 | 0.00E+00 | 0 | down | sense_intronic |
| NONRATT022363 | -Inf | 1.09E-02 | 0.37 | down | lincRNA |
| NONRATT022382 | -1.99 | 1.33E-02 | 0.42 | down | sense_overlapping |
| NONRATT022384 | -1.45 | 1.99E-03 | 0.11 | down | lincRNA |
| NONRATT023434 | -4.21 | 2.01E-02 | 0.54 | down | sense_overlapping |
| NONRATT023949 | -2.04 | 4.33E-02 | 0.82 | down | lincRNA |
| NONRATT024620 | -1.19 | 1.23E-03 | 0.08 | down | lincRNA |
| NONRATT024621 | -1.05 | 6.93E-03 | 0.27 | down | lincRNA |
| NONRATT024624 | -1.25 | 1.07E-03 | 0.07 | down | lincRNA |
| NONRATT024627 | -1.14 | 1.45E-03 | 0.08 | down | lincRNA |
| NONRATT024629 | -1.12 | 2.64E-03 | 0.13 | down | lincRNA |
| NONRATT024635 | -1.14 | 2.23E-02 | 0.58 | down | lincRNA |
| NONRATT024933 | -1.24 | 2.56E-02 | 0.63 | down | sense_overlapping |
| NONRATT024934 | -1.05 | 2.23E-02 | 0.58 | down | antisense |
| NONRATT025697 | -3.21 | 3.77E-02 | 0.77 | down | antisense |
| NONRATT025936 | -1.04 | 8.73E-03 | 0.31 | down | sense_intronic |
| NONRATT026602 | -1.11 | 4.61E-02 | 0.85 | down | sense_overlapping |
| NONRATT027353 | -3.49 | 3.77E-02 | 0.77 | down | sense_overlapping |
| NONRATT027644 | -1.51 | 1.74E-05 | 0.00 | down | sense_overlapping |
| NONRATT028105 | -3.84 | 2.01E-02 | 0.54 | down | sense_overlapping |
| NONRATT029020 | -4.43 | 2.01E-02 | 0.54 | down | sense_intronic |
| NONRATT029046 | -1.09 | 6.65E-05 | 0.01 | down | antisense |
| NONRATT029047 | -1.14 | 2.26E-03 | 0.12 | down | antisense |
| NONRATT029125 | -1.05 | 6.24E-03 | 0.25 | down | lincRNA |
| NONRATT029127 | -1.02 | 4.37E-03 | 0.19 | down | lincRNA |
| NONRATT029758 | -3.94 | 3.77E-02 | 0.77 | down | lincRNA |
| NONRATT031254 | -1.72 | 3.73E-02 | 0.77 | down | sense_overlapping |
| **L24/C** |  |  |  |  |  |
| NONRATT000533 | -1.12 | 4.01E-02 | 0.98 | down | sense_overlapping |
| NONRATT000963 | -1.28 | 9.59E-04 | 0.11 | down | lincRNA |
| NONRATT002754 | -1.43 | 4.51E-02 | 0.98 | down | lincRNA |
| NONRATT003494 | -3.71 | 7.11E-03 | 0.46 | down | sense_overlapping |
| NONRATT003737 | -1.45 | 2.76E-02 | 0.98 | down | sense_overlapping |
| NONRATT004080 | -2.57 | 1.29E-02 | 0.66 | down | lincRNA |
| NONRATT004138 | -2.13 | 1.29E-02 | 0.66 | down | sense_overlapping |
| NONRATT004410 | -5.58 | 1.87E-02 | 0.83 | down | sense_overlapping |
| NONRATT004414 | -2.95 | 4.29E-02 | 0.98 | down | sense_intronic |
| NONRATT004750 | -1.35 | 3.57E-02 | 0.98 | down | sense_overlapping |
| NONRATT005334 | -2.09 | 4.00E-03 | 0.31 | down | sense_overlapping |
| NONRATT005335 | -1.03 | 3.55E-02 | 0.98 | down | sense_overlapping |
| NONRATT005867 | -1.12 | 2.51E-03 | 0.22 | down | sense_intronic |
| NONRATT005952 | -1.05 | 1.65E-02 | 0.76 | down | antisense |
| NONRATT005954 | -1.17 | 3.10E-02 | 0.98 | down | sense_overlapping |
| NONRATT005957 | -1.76 | 1.67E-02 | 0.77 | down | sense_overlapping |
| NONRATT006395 | -3.77 | 3.94E-03 | 0.31 | down | lincRNA |
| NONRATT006743 | -1.84 | 2.26E-02 | 0.90 | down | lincRNA |
| NONRATT006758 | -2.07 | 3.96E-02 | 0.98 | down | sense_overlapping |
| NONRATT007528 | -1.23 | 6.54E-10 | 0.00 | down | sense_intronic |
| NONRATT007549 | -1.81 | 2.26E-02 | 0.90 | down | lincRNA |
| NONRATT007569 | -1.09 | 4.61E-02 | 0.98 | down | sense_overlapping |
| NONRATT007622 | -3.39 | 3.55E-02 | 0.98 | down | antisense |
| NONRATT007762 | -1.41 | 4.51E-02 | 0.98 | down | sense_overlapping |
| NONRATT008174 | -1.08 | 7.67E-03 | 0.48 | down | sense_intronic |
| NONRATT008234 | -2.39 | 4.29E-02 | 0.98 | down | antisense |
| NONRATT008642 | -1.78 | 1.99E-02 | 0.86 | down | lincRNA |
| NONRATT009350 | -2.65 | 4.29E-02 | 0.98 | down | sense_overlapping |
| NONRATT009377 | -1.94 | 0.00E+00 | 0 | down | lincRNA |
| NONRATT009510 | -1.31 | 4.51E-02 | 0.98 | down | sense_overlapping |
| NONRATT009710 | -2.39 | 4.29E-02 | 0.98 | down | antisense |
| NONRATT009762 | -1.03 | 2.74E-02 | 0.98 | down | sense_overlapping |
| NONRATT009800 | -1.02 | 2.74E-02 | 0.98 | down | sense_overlapping |
| NONRATT009823 | -1.54 | 3.34E-04 | 0.05 | down | lincRNA |
| NONRATT009835 | -1.79 | 1.99E-02 | 0.86 | down | lincRNA |
| NONRATT009895 | -3.39 | 3.55E-02 | 0.98 | down | sense_overlapping |
| NONRATT010410 | -1.04 | 4.89E-04 | 0.07 | down | sense_intronic |
| NONRATT011167 | -1.37 | 4.51E-02 | 0.98 | down | lincRNA |
| NONRATT011531 | -1.94 | 4.29E-02 | 0.98 | down | lincRNA |
| NONRATT011680 | -2.96 | 4.29E-02 | 0.98 | down | sense_overlapping |
| NONRATT011688 | -1.68 | 3.36E-02 | 0.98 | down | other |
| NONRATT012335 | -3.34 | 7.11E-03 | 0.46 | down | sense_overlapping |
| NONRATT012611 | -3.48 | 1.87E-02 | 0.83 | down | sense_overlapping |
| NONRATT012754 | -1.01 | 3.70E-05 | 0.01 | down | sense_overlapping |
| NONRATT012877 | -Inf | 3.55E-02 | 0.98 | down | lincRNA |
| NONRATT013954 | -1.23 | 1.36E-02 | 0.69 | down | sense_overlapping |
| NONRATT014771 | -1.53 | 4.51E-02 | 0.98 | down | sense_intronic |
| NONRATT016456 | -1.46 | 1.43E-04 | 0.03 | down | sense_overlapping |
| NONRATT017349 | -1.41 | 3.25E-12 | 0.00 | down | sense_overlapping |
| NONRATT017357 | -3.77 | 3.55E-02 | 0.98 | down | sense_overlapping |
| NONRATT017495 | -1.50 | 2.23E-02 | 0.90 | down | sense_overlapping |
| NONRATT017739 | -1.40 | 4.51E-02 | 0.98 | down | lincRNA |
| NONRATT018201 | -2.29 | 3.96E-02 | 0.98 | down | lincRNA |
| NONRATT018321 | -2.67 | 0.00E+00 | 0 | down | sense_overlapping |
| NONRATT018831 | -1.96 | 2.26E-02 | 0.90 | down | lincRNA |
| NONRATT019200 | -1.72 | 4.45E-04 | 0.07 | down | lincRNA |
| NONRATT019524 | -1.04 | 3.32E-03 | 0.28 | down | lincRNA |
| NONRATT020009 | -2.98 | 3.55E-02 | 0.98 | down | sense_overlapping |
| NONRATT020047 | -2.07 | 4.29E-02 | 0.98 | down | sense_overlapping |
| NONRATT021154 | -1.94 | 9.92E-04 | 0.11 | down | sense_intronic |
| NONRATT021330 | -4.07 | 3.55E-02 | 0.98 | down | sense_overlapping |
| NONRATT022337 | -1.94 | 0.00E+00 | 0 | down | sense_intronic |
| NONRATT022476 | -1.04 | 1.45E-02 | 0.71 | down | sense_overlapping |
| NONRATT022934 | -1.23 | 7.01E-03 | 0.46 | down | lincRNA |
| NONRATT023115 | -1.27 | 2.21E-02 | 0.89 | down | sense_overlapping |
| NONRATT024196 | -1.15 | 1.07E-02 | 0.60 | down | lincRNA |
| NONRATT025659 | -1.01 | 4.61E-02 | 0.98 | down | lincRNA |
| NONRATT025697 | -3.24 | 3.55E-02 | 0.98 | down | antisense |
| NONRATT027511 | -1.24 | 1.36E-02 | 0.69 | down | sense_overlapping |
| NONRATT027644 | -1.22 | 1.30E-04 | 0.03 | down | sense_overlapping |
| NONRATT027777 | -1.77 | 3.96E-02 | 0.98 | down | lincRNA |
| NONRATT029125 | -1.33 | 5.21E-04 | 0.07 | down | lincRNA |
| NONRATT029288 | -1.03 | 4.61E-02 | 0.98 | down | sense_overlapping |
| NONRATT029334 | -1.65 | 3.36E-02 | 0.98 | down | sense_overlapping |
| NONRATT029335 | -1.55 | 2.76E-02 | 0.98 | down | sense_overlapping |
| NONRATT029952 | -1.18 | 1.06E-03 | 0.11 | down | lincRNA |
| NONRATT030187 | -1.44 | 4.51E-02 | 0.98 | down | sense_overlapping |
| NONRATT031090 | -1.27 | 3.41E-02 | 0.98 | down | sense_overlapping |
| NONRATT031134 | -2.18 | 4.29E-02 | 0.98 | down | lincRNA |

### Characterization of the lncRNAs from RNA-seq data

The single-exon and multi-exon numbers occupied 10,012 (38.89%) and 15,732 (61.11%) of 25,745 lncRNA transcripts. There were 1,743 (8.23%) single exons and 19,439 (91.77%) multi-exons in the 21,183 mRNA transcripts (Fig. 6A). The length distribution analysis demonstrated that the lncRNAs and mRNAs mainly changed in the range >1,000 bp (Fig. 6B). Opening reading frame (ORF) analysis was based on the principle of six-frame translation, which aimed to provide evidence for gene prediction. From this prediction algorithm, it was clear that the ORFs coded by mRNA were longer than those coded by lncRNAs. It is possible that lncRNAs can code 500 amino acids, which is the classical hallmark of protein-coding genes (both from ncRNAs and mRNAs) (Fig. 6C and D).

**Fig 6:**
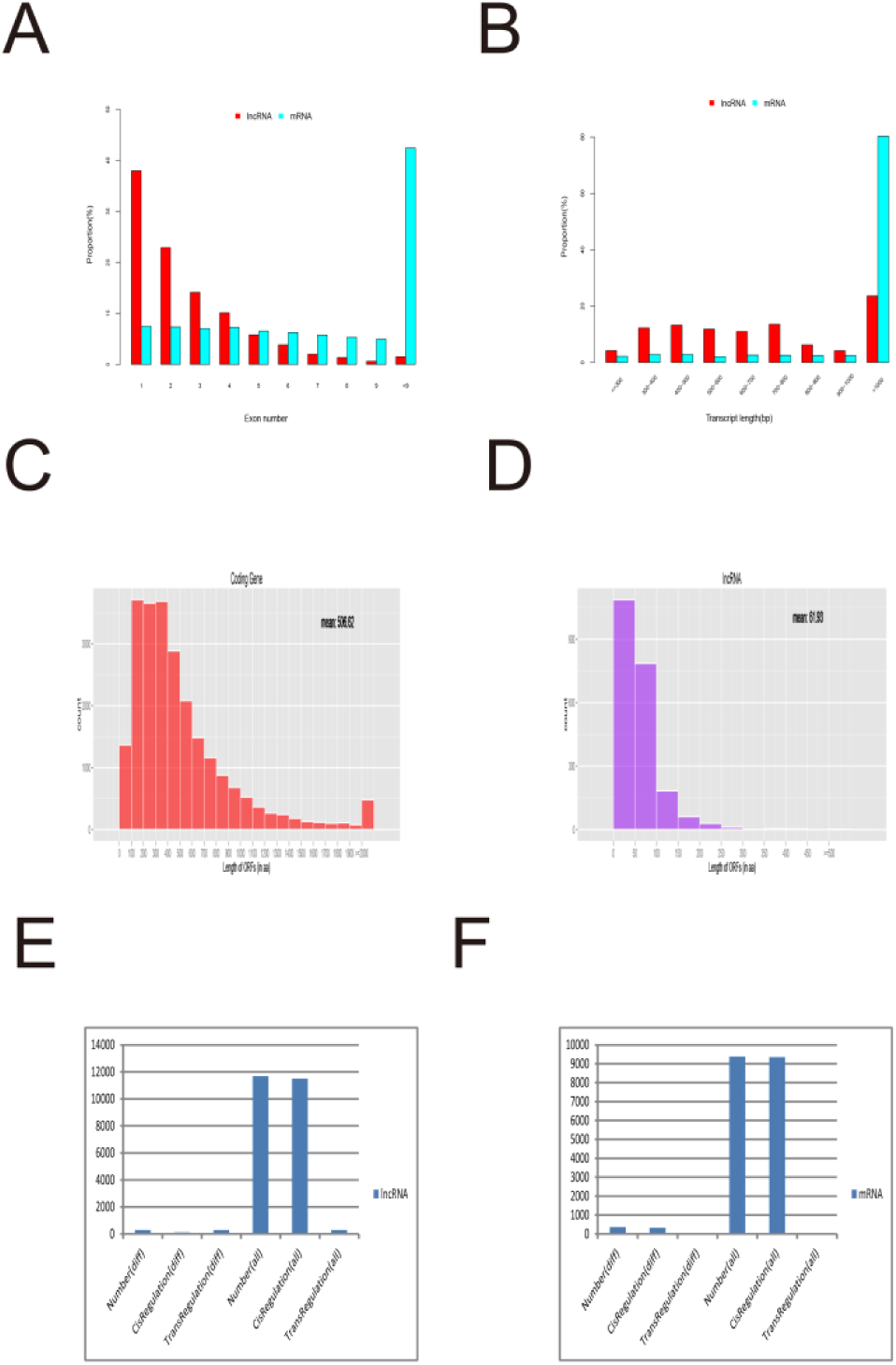
Detailed characteristics of significantly altered lncRNAs induced by LPS under different conditions in rodent brains. (A) Exon number of the transcripts of lncRNAs and mRNAs. (B) Length distribution of lncRNAs and mRNAs. (C) and (D) represent the length distribution of ORFs of lncRNAs and mRNAs. The x axis shows the length of the ORF (expressed by number of amino acids) and the y axis shows the number of ORFs in the IncRNA or mRNA sequence. (E) and (F) represent the regulation mode of expression of IncRNAs and mRNAs.

## Discussion

The mammalian brain expresses an abundant number of lncRNAs that have gradually become novel modulators for intervening in neuronal disease [16]. Using a rodent model of LPS-induced SAE and deep RNA sequencing (RNA-seq), we profiled LPS-modulated transcriptomes in adult rodent brains to determine the mechanism involved in neurological dysfunction after sepsis. In the present study, RNA-seq analysis of rodent brains under two conditions of LPS treatment revealed 195 responsive genes, among which, 38 mRNAs were changed in both groups. These results suggest that lncRNAs play critical roles in regulating gene expression, neurogenesis, function and disease in the CNS, which is in accordance with previous reports [17-20].

Various lncRNAs are highly expressed in the nervous system, which modulate gene expression via diverse mechanisms, including genetic imprinting, mRNA decay, chromatin remodeling, gene network reprogramming, and translational regulation [16, 21]. lncRNA dysfunction is related to the onset and progress of nervous system disorders [16, 22]. The growing evidence evokes awareness of the importance of lncRNAs in the CNS. However, because these molecules are novel and poorly understood, their role in defense against infection is largely unexplored. The prevalence of high-throughput RNA-seq provides an opportunity to map the lncRNA transcriptome. Comprehensive transcriptomic analysis helps us to understand the gene regulatory mechanisms in the LPS stress response in rats.

Recently, the pathogenesis of neural dysfunction and intellectual disorder is increasingly focused on defects in the microtubulin system [23]. Microtubules are an essential component of the cytoskeleton of almost all eukaryotic cells, and they are also responsible for the structural backbone of axons that participates in a diverse array of indispensable neural functions throughout life. They are enriched in the cell body and in the axonal initial and distal segments, such stable distribution of microtubules is vital to normal function of neurons [24]. If the process of microtubule production and polymerization in the CNS were changed, it would be reasonable to believe that there would also be morphogenetic changes in axons and many cellular processes, such as growth, division, polarity, migration, organelle positioning and cell-wall deposition [25-28]. Furthermore, these highly dynamic structures are intracellular transport trackers for signal transduction, organelle positioning and nutrient supplementation between cell bodies and processes [29, 30]. Their adaptive plasticity is strong, which can quickly respond to changes in the environment [24].

LPS is toxic to cultured eukaryotic cells and inhibites in vitro microtubule formation[31]. LPS targeted to intracellular microtubule architecture results in microtubule disassembly[32]. LPS can also bind to tubulin to cause microtubule fragmentation and shortening[33]. In the present study, after 6 or 24 h exposure to LPS, the comprehensive profiling suggested that protein-coding genes were frequently enriched in GO:007023 (post-chaperonin tubulin folding change). It is acknowledged that tubulins are the most highly conserved proteins across species [34]. The correct and complex folding of α- and β-tubulin is assisted by chaperonins, then tubulins are further processed to reach their final functional heterodimers, mediated by a set of five different tubulin binding cofactors, TBCA, TBCB, TBCC, TBCD and TBCE [35, 36]. In the present study, the most preferentially affected tubulin gene, *Tbcc,* which encodes one of the post-chaperonins, was significantly altered after 6 or 24 h exposure to LPS (*p* < 0.01). *Tbcc* is present at the centrosome, which is the organization center during the early phase of microtubule formation [37]. It is also involved in mitosis, which points to its vital role in eukaryotic genomic stability [37]. *Tbcc* is associated with the assembly of polarized α- and β-tubulin heterodimers in a head-to-tail fashion [38]. The ability to polymerize microtubules is a key cytoskeletal process in axonal growth and maintenance [37, 39].

Based on BLAST (http://www.ncbi.nlm.nih.gov/BLAST/) analysis, our newly identified lncRNAs (NONRATT020317) in rodent brains shared many characteristics with *BC1,* which implied that both possess analogous function in rodents. Rat *BC1* is a highly conserved ncRNA exclusively expressed in the rodent nervous system [40, 41], and its counterpart *BC200* is expressed in human brains [42, 43]. Both of them are located in postsynaptic domains of neurons. Furthermore, *BC1* and *BC200* are expressed in the same subpopulation of neurons in rats and primates, respectively [42]. Thus, the two kinds of RNA are probably functional analogs. *BC1* is formed in neurons and transferred through microtubules to synapses. It controls microtubule construction through inhibiting protein synthesis in dendrites, to modulate signal transduction [44, 45]. The mislocalization and overexpression of *BC1* result in microtubular malformation, and microtubules are closely associated with synaptic plasticity [46]. It is reported that *BC1* knockout mice have lower survival rates than control mice, which is manifested as more anxiety and lower exploratory behavior [47]. Dysfunction of primate-specific *BC200* impairs the delivery of RNA to the synapses, which is the main pathogenetic process of neurodegenerative diseases, such as Alzheimer’s disease [42]. Similarly, NONRATT020317 expressed in adult rat brains is probably responsible for modulating synaptic plasticity.

We found that the functional pathway KEGG 04360: axon guidance was significantly changed, which provides further evidence that microtubules are affected by inflammatory sequelae. Axon guidance guides the growth and development of axons in immature brains, whereas in developed brains, it controls synaptic plasticity [48, 49]. The aforementioned results verify that, after the inflammatory response, CNS dysfunction is partly associated with microtubular dysfunction and axon guidance. However, the exact mechanism of involvement of *BC1* in SAE remains to be elucidated. Elucidation of unknown expression patterns would be useful to explore the specific potential for this lncRNA in clinical translation.

The results of RNA-seq and the predictions of bioinformatics are preliminary, so evidence for lncRNAs in CNS dysfunction in sepsis is still needed. Although detailed functional validation was beyond the scope of this study, accurate verification of novel lncRNAs warrants further investigation through biological experiments. Additionally, with the help of biotechnological developments, the function of more lncRNAs in different types of cell stress will be established. Further studies could be conducted to determine the complex noncoding transcriptional networks in SAE. There is an urgency to develop genetic model systems and identify their functions *in vivo.* Finally, future research should be based on the prioritized lncRNAs to examine how the related noncoding transcriptional pathways intersect.

## Conclusions

This study is believed to be the first identification of LPS-responsive lncRNAs and mRNAs in adult rodent brains, with the potential to discover an unknown lncRNA-based molecular mechanism in SAE. The predicted lncRNA-mediated effects on microtubules could represent a mechanism for neural damage by LPS. For the preliminary characterization of the bioinformatics tool, the molecular and functional contributions are still warranted for further study by the integrational neural model. Undoubtedly, these data may also provide a basis for the new biomarkers, such as lncRNAs/mRNAs, with the aim of achieving early diagnosis or attenuating LPS-induced CNS sequelae.

## Abbreviations

CNS: Central nervous system
GO: Gene Ontology
ICU: Intensive care unit
KEGG: Kyoto Encyclopedia of Genes and Genomes
LPS: lipopolysaccharide
LncRNAs: Long non-coding RNAs
NT: Long non-coding RNAs
RPKM: Reads per kilobase per million mapped reads
SAE: Sepsis-associated encephalopathy
SD: Sprague-Dawley

## Author’s contributions

Wenchong Sun was a major contributor in manuscript writing and data analysis. Ling Pei gave an important direction for study design. Zuodi Liang participated in animal care and RNA extraction. All authors read and approved the final manuscript.

